## Supplemental Figs 1-5 for "The impact of primary colonizers on the community composition of river biofilm"

**Supplemental figures in support of our results include the following**.

1. Supplemental Fig 1. The top 25 populations of river water and river biofilm at 4-, 8- and 24-hours.
2. Supplemental Fig 2. Taxa S and Chao 1 diversity measures and evenness of river biofilms at 4-, 8- and 24-hours.
3. Supplemental Fig 3.  ANOSIM and PERMANOVER analysis of biofilm communities at 4-, 8- and 24-hours.
4. Supplemental Fig 4.  Bray-Curtis similarity derived from the sum of four replicates for each sample.
5. Supplemental Fig 5.  NMDS, ANOSIM and PERMANOVER analyses of biofilm communities at 4- & 8-hours.

**Supplemental Fig 1**

The Top 25 Populations of River Water and River Biofilm. Panels A, C & E are sorted according to the numerical abundance of river water populations. Panels B, D & F are sorted according to the numerical abundance of river biofilm populations. Panels A&B, C&D and E&F are samples at 4, 8 and 24 hours respectively. Populations that were isolated from eggs in previous studies are indicated with arrows.

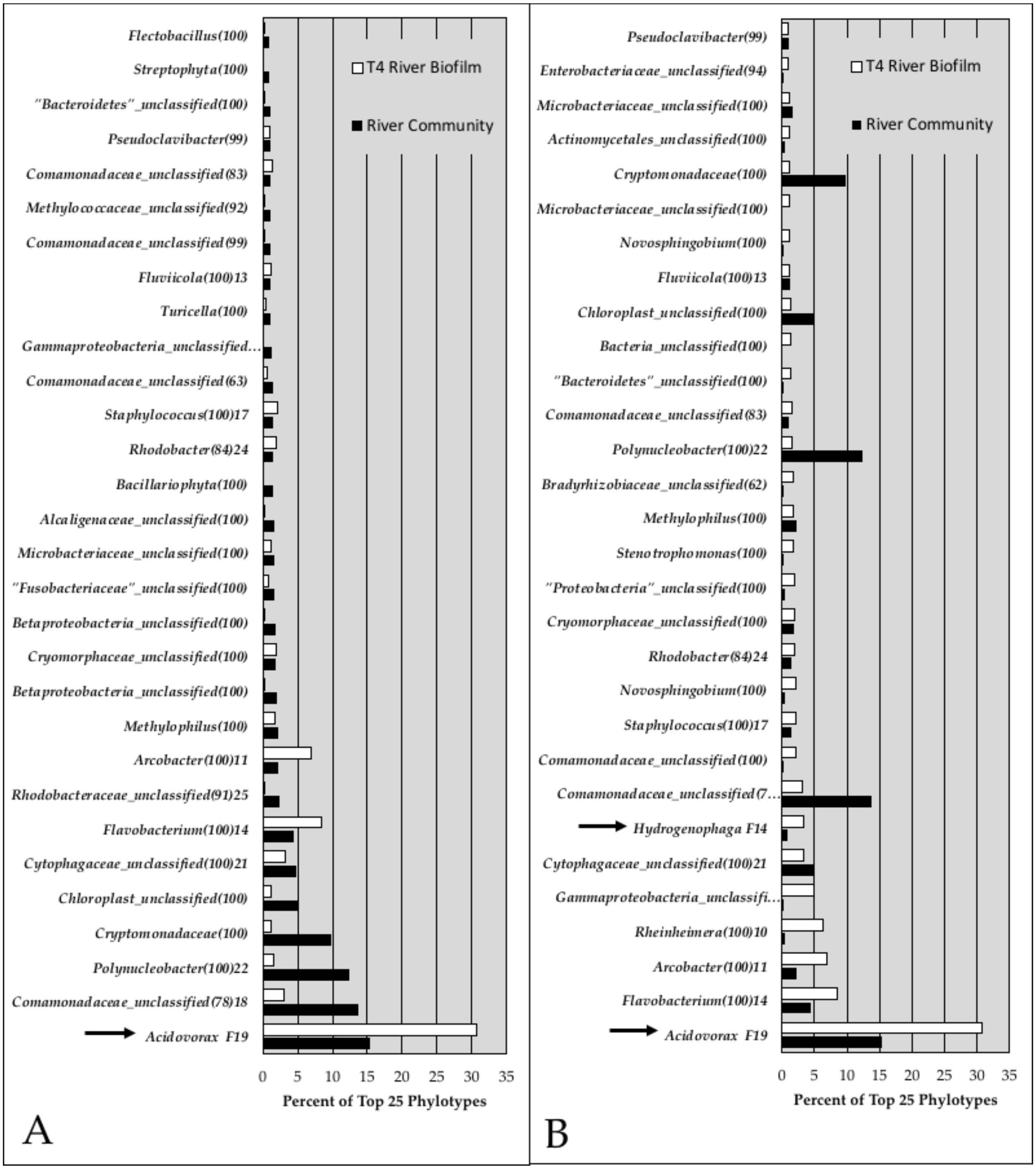

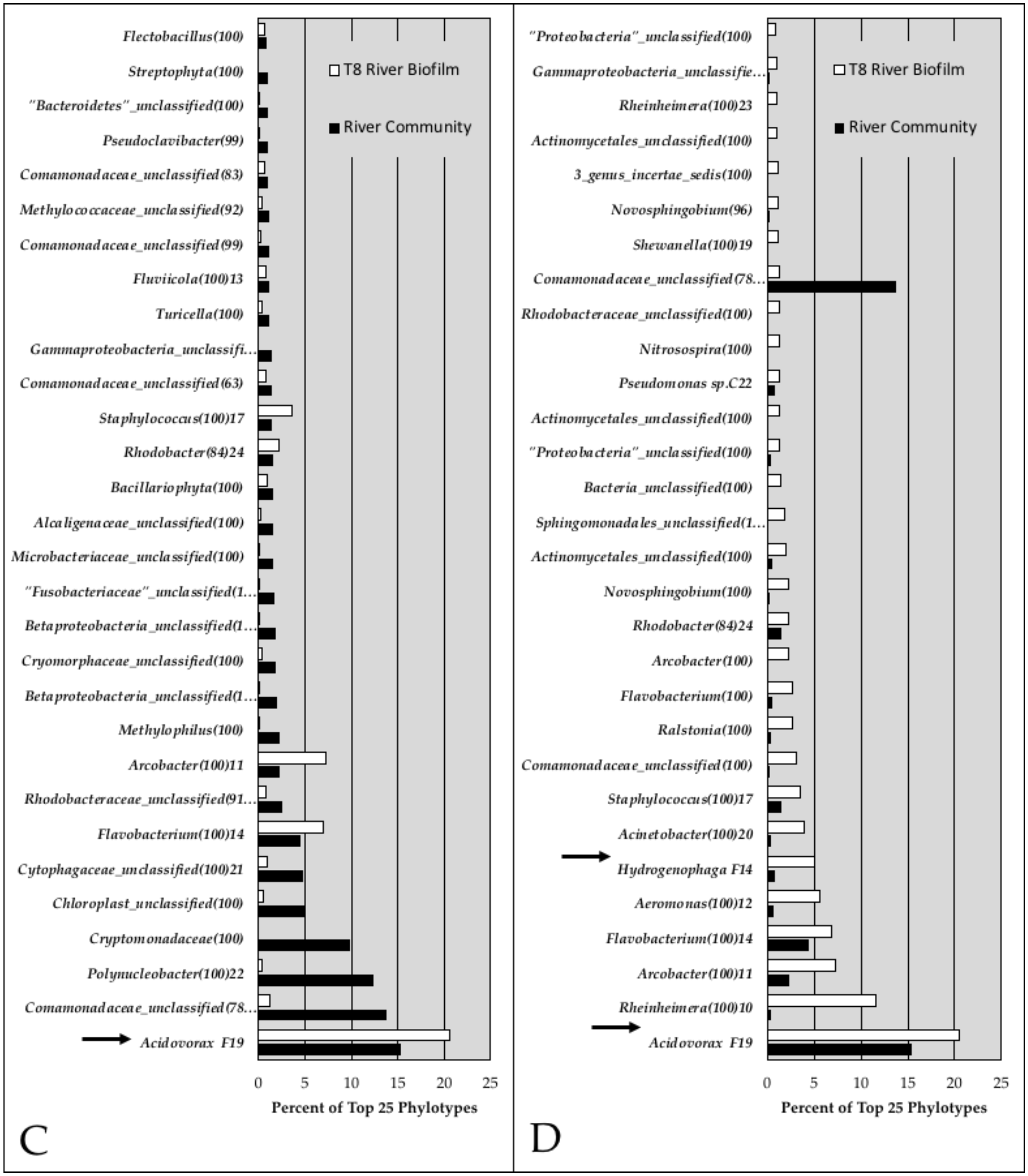

T8 River Biofilm
River Community
Flectobacillus(100)
Streptophyta(100)
"Bacteroidetes"_unclassified(100)
Pseudoclavibacter(99)
Comamonadaceae_unclassified(83)
Methylococcaceae_unclassified(92)
Comamonadaceae_unclassified(99)
Fluviicola(100)13
Turicella(100)
Gammaproteobacteria_unclassifi...
Comamonadaceae_unclassified(63)
Staphylococcus(100)17
Rhodobacter(84)24
Bacillariophyta(100)
Alcaligenaceae_unclassified(100)
Microbacteriaceae_unclassified(100)
"Fusobacteriaceae"_unclassified(1...
Betaproteobacteria_unclassified(1...
Cryomorphaceae_unclassified(100)
Betaproteobacteria_unclassified(1...
Methylophilus(100)
Arcobacter(100)11
Rhodobacteraceae_unclassified(91...
Flavobacterium(100)14
Cytophagaceae_unclassified(100)21
Chloroplast_unclassified(100)
Cryptomonadaceae(100)
Polynucleobacter(100)22
Comamonadaceae_unclassified(78...
Acidovorax F19
0 5 10 15 20 25
Percent of Top 25 Phylotypes
C
"Proteobacteria"_unclassified(100)
Gammaproteobacteria_unclassifie...
Rheinheimera(100)23
Actinomycetales_unclassified(100)
3_genus_incertae_sedis(100)
Novosphingobium(96)
Shewanella(100)19
Comamonadaceae_unclassified(78...
Rhodobacteraceae_unclassified(100)
Nitrosospira(100)
Pseudomonas sp.C22
Actinomycetales_unclassified(100)
"Proteobacteria"_unclassified(100)
Bacteria_unclassified(100)
Sphingomonadales_unclassified(1...
Actinomycetales_unclassified(100)
Novosphingobium(100)
Rhodobacter(84)24
Arcobacter(100)
Flavobacterium(100)
Ralstonia(100)
Comamonadaceae_unclassified(100)
Staphylococcus(100)17
Acinetobacter(100)20
Hydrogenophaga F14
Aeromonas(100)12
Flavobacterium(100)14
Arcobacter(100)11
Rheinheimera(100)10
Acidovorax F19
0 5 10 15 20 25
Percent of Top 25 Phylotypes
D

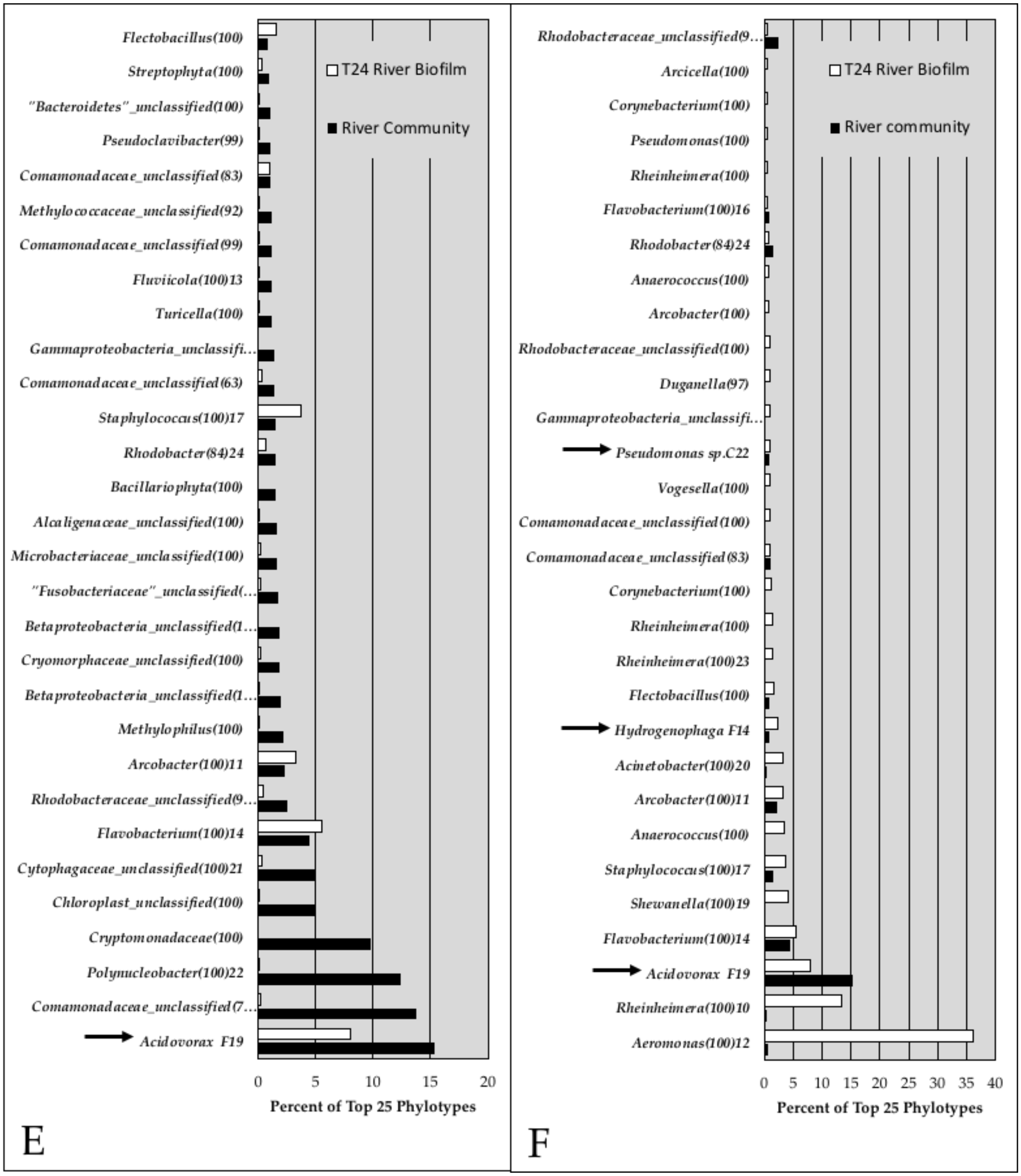

Flectobacillus(100)
Streptophyta(100)
"Bacteroidetes"_unclassified(100)
Pseudoclavibacter(99)
Comamonadaceae_unclassified(83)
Methylococcaceae_unclassified(92)
Comamonadaceae_unclassified(99)
Fluviicola(100)13
Turicella(100)
Gammaproteobacteria_unclassifi…
Comamonadaceae_unclassified(63)
Staphylococcus(100)17
Rhodobacter(84)24
Bacillariophyta(100)
Alcaligenaceae_unclassified(100)
Microbacteriaceae_unclassified(100)
"Fusobacteriaceae"_unclassified(…
Betaproteobacteria_unclassified(1…
Cryomorphaceae_unclassified(100)
Betaproteobacteria_unclassified(1…
Methylophilus(100)
Arcobacter(100)11
Rhodobacteraceae_unclassified(9…
Flavobacterium(100)14
Cytophagaceae_unclassified(100)21
Chloroplast_unclassified(100)
Cryptomonadaceae(100)
Polynucleobacter(100)22
Comamonadaceae_unclassified(7…
Acidovorax F19
T24 River Biofilm
River Community
0 5 10 15 20
Percent of Top 25 Phylotypes
E
Rhodobacteraceae_unclassified(9…
Arcicella(100)
Corynebacterium(100)
Pseudomonas(100)
Rheinheimera(100)
Flavobacterium(100)16
Rhodobacter(84)24
Anaerococcus(100)
Arcobacter(100)
Rhodobacteraceae_unclassified(100)
Duganella(97)
Gammaproteobacteria_unclassifi…
Pseudomonas sp.C22
Vogesella(100)
Comamonadaceae_unclassified(100)
Comamonadaceae_unclassified(83)
Corynebacterium(100)
Rheinheimera(100)
Rheinheimera(100)23
Flectobacillus(100)
Hydrogenophaga F14
Acinetobacter(100)20
Arcobacter(100)11
Anaerococcus(100)
Staphylococcus(100)17
Shewanella(100)19
Flavobacterium(100)14
Acidovorax F19
Rheinheimera(100)10
Aeromonas(100)12
T24 River Biofilm
River community
0 5 10 15 20 25 30 35 40
Percent of Top 25 Phylotypes
F

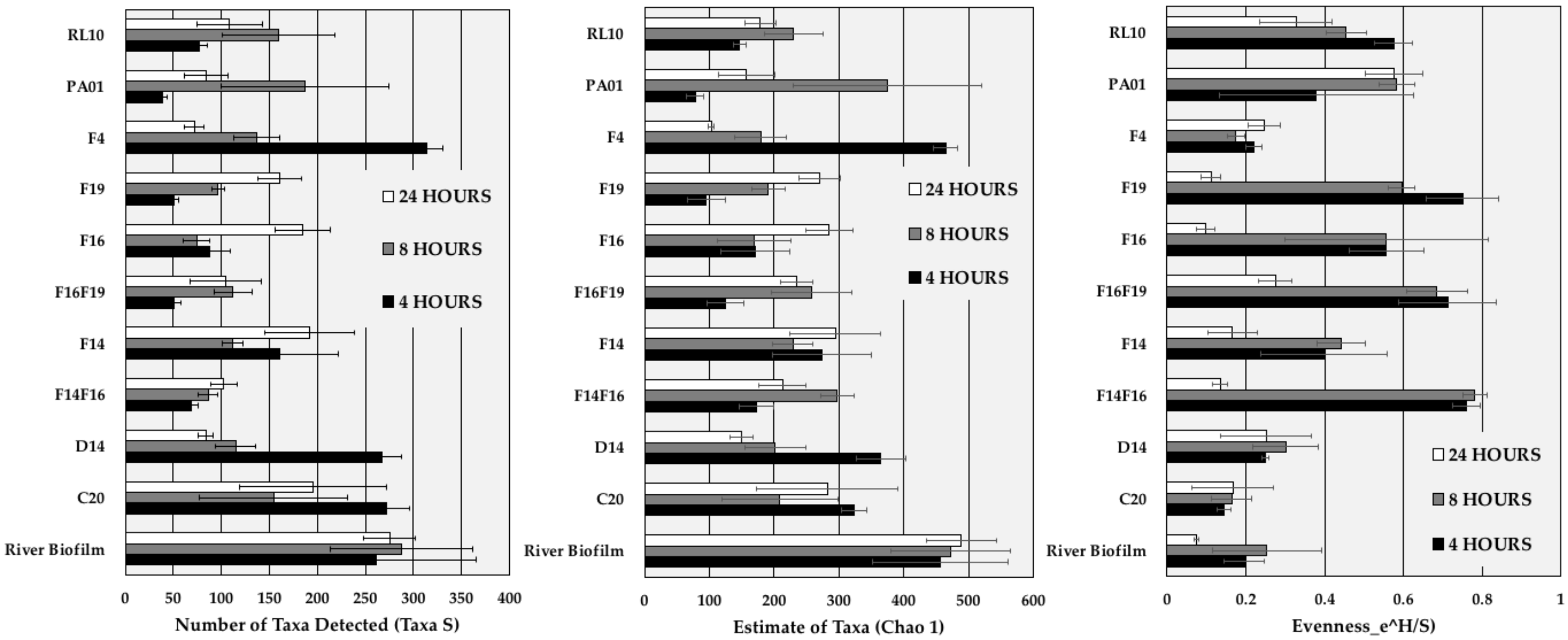


**Supplemental Fig 2.** Taxa S and Chao 1 diversity measures and Evenness of river biofilms at 4, 8 and 24 hours in the presence and absence of preestablished isolate biofilms. These measurements are derived from community analyses from which the founding isolate populations have been subtracted, thereby providing values that are not substantially influenced by the large population size of the established biofilms.

**Supplemental Fig 3**. ANOSIM and PERMANOVER analysis of biofilm communities at 4, 8 and 24 hours after removal of founding populations.

| **ANOSIM 4 hours** | **River Biofilm** | **C20** | **D14** | **F14-F16** | **F14** | **F16-F19** | **F16** | **F19** | **F4** | **PA01** | **RL10** | |
|---|---|---|---|---|---|---|---|---|---|---|---|---|
| **River Biofilm** | | 0.03 | 0.0309 | 0.0278 | 0.0302 | 0.0275 | 0.0288 | 0.0267 | 0.0295 | 0.0263 | 0.0264 | |
| **C20** | **0.5** | | 0.0295 | 0.0287 | 0.0257 | 0.0311 | 0.0265 | 0.0283 | 0.0302 | 0.0298 | 0.03 | |
| **D14** | **1** | **1** | | 0.0294 | 0.0294 | 0.0258 | 0.0277 | 0.0266 | 0.0255 | 0.0258 | 0.0287 | |
| **F14-F16** | **1** | **1** | **1** | | 0.026 | 0.0293 | 0.0291 | 0.0289 | 0.0291 | 0.0266 | 0.0273 | |
| **F14** | **1** | **1** | **0.7083** | **1** | | 0.031 | 0.0268 | 0.0284 | 0.0276 | 0.027 | 0.0288 | |
| **F16-F19** | **1** | **1** | **1** | **1** | **1** | | 0.0281 | 0.0312 | 0.0274 | 0.0307 | 0.027 | |
| **F16** | **1** | **1** | **0.9896** | **0.9375** | **0.625** | **0.9074** | | 0.026 | 0.0271 | 0.0286 | 0.0277 | |
| **F19** | **1** | **1** | **1** | **1** | **1** | **1** | **1** | | 0.0254 | 0.0303 | 0.0306 | |
| **F4** | **1** | **1** | **0.9583** | **1** | **0.7083** | **1** | **1** | **1** | | 0.029 | 0.0278 | |
| **PA01** | **1** | **1** | **1** | **0.9375** | **0.9167** | **1** | **0.9896** | **1** | **1** | | 0.0269 | **R value** |
| **RL10** | **1** | **1** | **1** | **1** | **1** | **1** | **0.9792** | **1** | **1** | **0.9688** | | P value |

| **PERMANOVER 4 hours** | **River Biofilm** | **C20** | **D14** | **F14-F16** | **F14** | **F16-F19** | **F16** | **F19** | **F4** | **PA01** | **RL10** | |
|---|---|---|---|---|---|---|---|---|---|---|---|---|
| **River Biofilm** | | 0.0324 | 0.0299 | 0.0287 | 0.0263 | 0.0302 | 0.0284 | 0.0267 | 0.0283 | 0.029 | 0.0289 | |
| **C20** | **2.896** | | 0.0315 | 0.0326 | 0.0294 | 0.0265 | 0.0272 | 0.0293 | 0.0266 | 0.0291 | 0.0311 | |
| **D14** | **8.672** | **45.66** | | 0.0271 | 0.0258 | 0.0296 | 0.0292 | 0.0276 | 0.0301 | 0.0281 | 0.0271 | |
| **F14-F16** | **12.21** | **79.72** | **60.09** | | 0.0287 | 0.0252 | 0.0303 | 0.0291 | 0.0284 | 0.0296 | 0.0277 | |
| **F14** | **11.55** | **62.78** | **14.82** | **1.729** | | 0.1211 | 0.0302 | 0.027 | 0.0245 | 0.4624 | 0.0285 | |
| **F16-F19** | **8.759** | **57.29** | **43.99** | **2.939** | **1.348** | | 0.0892 | 0.0814 | 0.0263 | 0.4508 | 0.0297 | |
| **F16** | **11.96** | **76.84** | **49.98** | **2.878** | **1.467** | **2.634** | | 0.0301 | 0.0268 | 0.0302 | 0.0296 | |
| **F19** | **12.26** | **80.23** | **62.1** | **4.518** | **1.776** | **1.846** | **3.908** | | 0.0309 | 0.0289 | 0.0291 | |
| **F4** | **8.47** | **38.89** | **6.934** | **65.04** | **17.22** | **47.34** | **54.34** | **66.66** | | 0.0303 | 0.0299 | |
| **PA01** | **12.18** | **73** | **29.76** | **1.257** | **1.05** | **0.9777** | **1.477** | **1.273** | **32.93** | | 0.0281 | **F value** |
| **RL10** | **12.17** | **79.08** | **56.95** | **7.234** | **1.681** | **5.649** | **4.458** | **8.449** | **62.08** | **1.432** | | P valiue |

| ANOSIM 8 hours | River Biofilm | C20 | D14 | F14-F16 | F14 | F16-F19 | F16 | F19 | F4 | PA01 | RL10 | |
|---|---|---|---|---|---|---|---|---|---|---|---|---|
| **River Biofilm** | | 0.1393 | 0.0316 | 0.0307 | 0.0285 | 0.0284 | 0.0281 | 0.0288 | 0.0313 | 0.0284 | 0.0314 | |
| **C20** | **0.1146** | | 0.4915 | 0.0292 | 0.0287 | 0.0334 | 0.0275 | 0.0296 | 0.0298 | 0.0272 | 0.0273 | |
| **D14** | **0.5208** | **0.03125** | | 0.0249 | 0.0293 | 0.0324 | 0.0282 | 0.0266 | 0.029 | 0.0279 | 0.0298 | |
| **F14-F16** | **1** | **1** | **1** | | 0.0275 | 0.0318 | 0.0272 | 0.0284 | 0.0326 | 0.0313 | 0.0301 | |
| **F14** | **1** | **0.8854** | **1** | **1** | | 0.026 | 0.0305 | 0.0314 | 0.0298 | 0.032 | 0.0301 | |
| **F16-F19** | **1** | **1** | **1** | **0.9896** | **0.9063** | | 0.0297 | 0.0281 | 0.0293 | 0.0277 | 0.0291 | |
| **F16** | **1** | **0.9063** | **0.9479** | **0.8021** | **0.9688** | **0.9375** | | 0.0278 | 0.0287 | 0.0274 | 0.0266 | |
| **F19** | **1** | **1** | **1** | **1** | **0.9896** | **0.6979** | **1** | | 0.0298 | 0.0294 | 0.0261 | |
| **F4** | **0.5** | **0.3958** | **0.4167** | **1** | **1** | **1** | **1** | **1** | | 0.0276 | 0.0276 | |
| **PA01** | **1** | **1** | **1** | **0.9479** | **0.8542** | **0.8854** | **0.7396** | **0.9583** | **1** | | 0.0296 | **R value** |
| **RL10** | **1** | **0.7917** | **0.8958** | **1** | **0.5833** | **0.8333** | **0.7708** | **0.7708** | **0.9167** | **0.75** | | P value |

| PERMANOVER 8 hours | River Biofilm | C20 | D14 | F14-F16 | F14 | F16-F19 | F16 | F19 | F4 | PA01 | RL10 | |
|---|---|---|---|---|---|---|---|---|---|---|---|---|
| **River Biofilm** | | 0.2614 | 0.0299 | 0.029 | 0.0287 | 0.0315 | 0.0285 | 0.0264 | 0.0326 | 0.029 | 0.0294 | |
| **C20** | **1.233** | | 0.2278 | 0.0298 | 0.0266 | 0.0284 | 0.028 | 0.0295 | 0.0288 | 0.0269 | 0.028 | |
| **D14** | **1.999** | **1.315** | | 0.0277 | 0.0283 | 0.0304 | 0.0252 | 0.0293 | 0.0277 | 0.0276 | 0.0286 | |
| **F14-F16** | **4.514** | **3.892** | **3.058** | | 0.0278 | 0.0289 | 0.0311 | 0.0301 | 0.0246 | 0.0276 | 0.0286 | |
| **F14** | **5.262** | **4.225** | **3.078** | **3.331** | | 0.028 | 0.0303 | 0.0297 | 0.0281 | 0.0303 | 0.0278 | |
| **F16-F19** | **3.711** | **3.154** | **2.429** | **2.288** | **2.658** | | 0.0311 | 0.03 | 0.0261 | 0.0324 | 0.0304 | |
| **F16** | **4.574** | **3.787** | **2.855** | **2.079** | **3.234** | **2.253** | | 0.0299 | 0.0273 | 0.0258 | 0.0292 | |
| **F19** | **5.142** | **4.323** | **3.305** | **3.618** | **2.906** | **1.961** | **3.397** | | 0.0269 | 0.0269 | 0.0272 | |
| **F4** | **3.37** | **2.376** | **3.661** | **7.92** | **9.504** | **6.239** | **7.94** | **9.264** | | 0.0267 | 0.0295 | |
| **PA01** | **3.344** | **2.918** | **2.226** | **2.407** | **2.974** | **1.95** | **2.242** | **2.749** | **5.674** | | 0.0263 | **F value** |
| **RL10** | **3.483** | **2.803** | **1.909** | **2.85** | **2.001** | **1.905** | **2.325** | **2.501** | **5.713** | **2.04** | | P valiue |

| ANOSIM 24 hours | River Biofilm | C20 | D14 | F14-F16 | F14 | F16-F19 | F16 | F19 | F4 | PA01 | RL10 | |
|---|---|---|---|---|---|---|---|---|---|---|---|---|
| **River Biofilm** | | 0.0279 | 0.0299 | 0.0267 | 0.0289 | 0.0278 | 0.0284 | 0.031 | 0.0283 | 0.0284 | 0.0276 | |
| **C20** | **1** | | 0.0586 | 0.0308 | 0.0278 | 0.0288 | 0.0262 | 0.0288 | 0.0842 | 0.0319 | 0.0582 | |
| **D14** | **1** | **0.4479** | | 0.0318 | 0.1102 | 0.0231 | 0.0281 | 0.0284 | 0.0301 | 0.0286 | 0.8599 | |
| **F14-F16** | **1** | **0.75** | **0.375** | | 0.0281 | 0.1137 | 0.0263 | 0.0287 | 0.0271 | 0.0292 | 0.0259 | |
| **F14** | **0.9896** | **0.6146** | **0.2083** | **0.5833** | | 0.0557 | 0.0259 | 0.1176 | 0.0297 | 0.2023 | 0.1433 | |
| **F16-F19** | **1** | **0.75** | **0.2917** | **0.1042** | **0.4167** | | 0.0281 | 0.0289 | 0.0268 | 0.0287 | 0.0586 | |
| **F16** | **0.9375** | **0.3854** | **0.4063** | **0.5** | **0.625** | **0.5313** | | 0.026 | 0.0291 | 0.0302 | 0.0279 | |
| **F19** | **1** | **0.625** | **0.4271** | **1** | **0.1771** | **0.8958** | **0.6146** | | 0.03 | 0.085 | 0.028 | |
| **F4** | **0.6146** | **0.2396** | **0.5** | **0.625** | **0.3229** | **0.625** | **0.25** | **0.3958** | | 0.0287 | 0.0882 | |
| **PA01** | **1** | **0.6771** | **0.25** | **0.7813** | **0.1563** | **0.4479** | **0.625** | **0.3125** | **0.5521** | | 0.0841 | **R value** |
| **RL10** | **1** | **0.5** | **-0.1667** | **0.6481** | **0.2593** | **0.3889** | **0.4815** | **0.5926** | **0.3333** | **0.2407** | | P value |

| PERMANOVER 24 hours | River Biofilm | C20 | D14 | F14-F16 | F14 | F16-F19 | F16 | F19 | F4 | PA01 | RL10 | |
|---|---|---|---|---|---|---|---|---|---|---|---|---|
| **River Biofilm** | | 0.0292 | 0.0311 | 0.0294 | 0.0253 | 0.0282 | 0.033 | 0.027 | 0.0273 | 0.0251 | 0.0289 | |
| **C20** | **10.92** | | 0.0598 | 0.0296 | 0.0271 | 0.0273 | 0.0337 | 0.0303 | 0.0878 | 0.0276 | 0.032 | |
| **D14** | **16.47** | **3.701** | | 0.0289 | 0.0312 | 0.0292 | 0.0277 | 0.0294 | 0.0282 | 0.0266 | 0.7359 | |
| **F14-F16** | **30.44** | **9.542** | **2.946** | | 0.026 | 0.0864 | 0.0295 | 0.0274 | 0.0305 | 0.0306 | 0.0294 | |
| **F14** | **16.51** | **5.861** | **2.748** | **6.153** | | 0.0281 | 0.0298 | 0.2247 | 0.0294 | 0.1449 | 0.1455 | |
| **F16-F19** | **28.55** | **9.098** | **2.529** | **1.637** | **4.737** | | 0.0306 | 0.0289 | 0.0304 | 0.0299 | 0.0299 | |
| **F16** | **12.35** | **4.56** | **2.567** | **2.535** | **3.37** | **2.99** | | 0.0304 | 0.0271 | 0.0302 | 0.0282 | |
| **F19** | **26.58** | **7.627** | **3.351** | **24.95** | **1.299** | **10.55** | **3.798** | | 0.0284 | 0.1105 | 0.0263 | |
| **F4** | **5.266** | **1.951** | **3.513** | **5.552** | **3.251** | **5.448** | **2.957** | **4.246** | | 0.0257 | 0.0283 | |
| **PA01** | **27.49** | **8.108** | **2.507** | **9.464** | **2.353** | **3.983** | **3.888** | **2.509** | **4.937** | | 0.146 | **F value** |
| **RL10** | **15.97** | **3.71** | **0.5149** | **4.033** | **1.703** | **2.737** | **2.308** | **2.55** | **2.788** | **1.686** | | P valiue |

**Supplemental Fig 4**. Bray-Curtis similarity derived from the sum of four replicates for each sample.

| | RedCedar | T4_RC_Biofilm | T8_RC_Biofilm | T24_RC_Biofilm | T4_C20 | T8_C20 | T24_C20 | T4_D14 | T8_D14 | T24_D14 | T4_F14 | T8_F14 | T24_F14 | T4_F14F16 | T8_F14F16 | T24_F14F16 | T4_F16 |
|---|---|---|---|---|---|---|---|---|---|---|---|---|---|---|---|---|---|
| **RedCedar** | 1 | 0.42814511 | 0.34239993 | 0.23528914 | 0.36706713 | 0.29309279 | 0.32604208 | 0.23096924 | 0.22380413 | 0.2228612 | 0.059939302 | 0.053469787 | 0.14513869 | 0.010056593 | 0.00828386 | 0.06538639 | 0.027408524 |
| **T4_RC_Biofilm** | 0.42814511 | 1 | 0.49881519 | 0.35584096 | 0.50792942 | 0.40667375 | 0.52596122 | 0.24055976 | 0.25701066 | 0.27027027 | 0.06078197 | 0.062571521 | 0.20459762 | 0.011505273 | 0.009278052 | 0.083640398 | 0.027999558 |
| **T8_RC_Biofilm** | 0.34239993 | 0.49881519 | 1 | 0.45094078 | 0.48634091 | 0.41930288 | 0.49512876 | 0.21131903 | 0.25891095 | 0.2634016 | 0.060368014 | 0.064701436 | 0.32569216 | 0.01030353 | 0.009190889 | 0.075850044 | 0.026766951 |
| **T24_RC_Biofilm** | 0.23528914 | 0.35584096 | 0.45094078 | 1 | 0.33199887 | 0.31864538 | 0.42623182 | 0.18299758 | 0.24032783 | 0.22272261 | 0.052572014 | 0.056815727 | 0.36817302 | 0.008745315 | 0.007855039 | 0.081303391 | 0.024076123 |
| **T4_C20** | 0.36706713 | 0.50792942 | 0.48634091 | 0.33199887 | 1 | 0.68357915 | 0.49684904 | 0.40641437 | 0.31768972 | 0.39951858 | 0.12436129 | 0.12063773 | 0.27269776 | 0.0231278 | 0.017248004 | 0.13225984 | 0.054619554 |
| **T8_C20** | 0.29309279 | 0.40667375 | 0.41930288 | 0.31864538 | 0.68357915 | 1 | 0.416357 | 0.41464538 | 0.36868762 | 0.42979617 | 0.12579978 | 0.14198311 | 0.24346734 | 0.026978319 | 0.021094049 | 0.13684211 | 0.065213899 |
| **T24_C20** | 0.32604208 | 0.52596122 | 0.49512876 | 0.42623182 | 0.49684904 | 0.416357 | 1 | 0.22963023 | 0.22819264 | 0.27586342 | 0.060790274 | 0.062536433 | 0.30194722 | 0.0107192 | 0.008259074 | 0.10354942 | 0.029339268 |
| **T4_D14** | 0.23096924 | 0.24055976 | 0.21131903 | 0.18299758 | 0.40641437 | 0.41464538 | 0.22963023 | 1 | 0.39739521 | 0.37483209 | 0.26973563 | 0.26077649 | 0.25344673 | 0.061287028 | 0.044323219 | 0.25110803 | 0.14253024 |
| **T8_D14** | 0.22380413 | 0.25701066 | 0.25891095 | 0.24032783 | 0.31768972 | 0.36868762 | 0.22819264 | 0.39739521 | 1 | 0.28702789 | 0.1162753 | 0.13378559 | 0.19402935 | 0.024337087 | 0.023021931 | 0.13049171 | 0.059109216 |
| **T24_D14** | 0.2228612 | 0.27027027 | 0.2634016 | 0.22272261 | 0.39951858 | 0.42979617 | 0.27586342 | 0.37483209 | 0.28702789 | 1 | 0.12770916 | 0.14190723 | 0.21971716 | 0.023743788 | 0.021743136 | 0.17423462 | 0.069978632 |
| **T4_F14** | 0.059939302 | 0.06078197 | 0.060368014 | 0.052572014 | 0.12436129 | 0.12579978 | 0.060790274 | 0.26973563 | 0.1162753 | 0.12770916 | 1 | 0.46001322 | 0.16275148 | 0.18613733 | 0.14831754 | 0.1932889 | 0.29485315 |
| **T8_F14** | 0.053469787 | 0.062571521 | 0.064701436 | 0.056815727 | 0.12063773 | 0.14198311 | 0.062536433 | 0.26077649 | 0.13378559 | 0.14190723 | 0.46001322 | 1 | 0.16163377 | 0.19905213 | 0.1984127 | 0.21835559 | 0.26017943 |
| **T24_F14** | 0.14513869 | 0.20459762 | 0.32569216 | 0.36817302 | 0.27269776 | 0.24346734 | 0.30194722 | 0.25344673 | 0.19402935 | 0.21971716 | 0.16275148 | 0.16163377 | 1 | 0.03096951 | 0.028252074 | 0.25138449 | 0.053205 |
| **T4_F14F16** | 0.010056593 | 0.011505273 | 0.01030353 | 0.008745315 | 0.0231278 | 0.026978319 | 0.0107192 | 0.061287028 | 0.024337087 | 0.023743788 | 0.18613733 | 0.19905213 | 0.03096951 | 1 | 0.42314991 | 0.074981329 | 0.35055866 |
| **T8_F14F16** | 0.00828386 | 0.009278052 | 0.009190889 | 0.007855039 | 0.017248004 | 0.021094049 | 0.008259074 | 0.044323219 | 0.023021931 | 0.021743136 | 0.14831754 | 0.1984127 | 0.028252074 | 0.42314991 | 1 | 0.068532096 | 0.27323944 |
| **T24_F14F16** | 0.06538639 | 0.083640398 | 0.075850044 | 0.081303391 | 0.13225984 | 0.13684211 | 0.10354942 | 0.25110803 | 0.13049171 | 0.17423462 | 0.1932889 | 0.21835559 | 0.25138449 | 0.074981329 | 0.068532096 | 1 | 0.096586886 |
| **T4_F16** | 0.027408524 | 0.027999558 | 0.026766951 | 0.024076123 | 0.054619554 | 0.065213899 | 0.029339268 | 0.14253024 | 0.059109216 | 0.069978632 | 0.29485315 | 0.26017943 | 0.053205 | 0.35055866 | 0.27323944 | 0.096586886 | 1 |
| **T8_F16** | 0.029446269 | 0.032862191 | 0.033355423 | 0.032040658 | 0.05958267 | 0.07384484 | 0.033498996 | 0.1540792 | 0.084842764 | 0.11082588 | 0.25913242 | 0.25312184 | 0.081991215 | 0.26853707 | 0.25319865 | 0.08195341 | 0.39076758 |
| **T24_F16** | 0.17878928 | 0.2673833 | 0.28539028 | 0.28857627 | 0.32755506 | 0.32103838 | 0.32213641 | 0.29254518 | 0.25195179 | 0.27187786 | 0.087854902 | 0.087523932 | 0.29657956 | 0.018485514 | 0.016255443 | 0.34393064 | 0.051116646 |
| **T4_F16F19** | 0.004512171 | 0.005641875 | 0.006135794 | 0.004928205 | 0.011181834 | 0.013349515 | 0.005237887 | 0.030134446 | 0.014080196 | 0.0168492 | 0.06366782 | 0.072371222 | 0.010824313 | 0.22197055 | 0.22043628 | 0.023955774 | 0.21777422 |
| **T8_F16F19** | 0.022584839 | 0.022068966 | 0.022511771 | 0.018050383 | 0.051358147 | 0.047029315 | 0.02149265 | 0.097748262 | 0.05050738 | 0.047382826 | 0.21903052 | 0.21024419 | 0.05901044 | 0.18710359 | 0.17340426 | 0.094136418 | 0.23117803 |
| **T24_F16F19** | 0.10741389 | 0.11466646 | 0.11866025 | 0.094321242 | 0.1596047 | 0.2005418 | 0.14488253 | 0.25483476 | 0.24019628 | 0.22951889 | 0.13450292 | 0.16976444 | 0.16859519 | 0.032809773 | 0.037982058 | 0.4451631 | 0.067626381 |
| **T4_F19** | 0.007782716 | 0.00764058 | 0.008670197 | 0.006258941 | 0.015847861 | 0.019206497 | 0.006967798 | 0.045626505 | 0.018989653 | 0.021154203 | 0.10653266 | 0.10883797 | 0.016405873 | 0.18200409 | 0.1573499 | 0.022097775 | 0.21130952 |
| **T8_F19** | 0.025543033 | 0.023160583 | 0.026953228 | 0.023790118 | 0.050066644 | 0.058991739 | 0.026098967 | 0.12766411 | 0.060967761 | 0.073258152 | 0.26494789 | 0.30466989 | 0.066060127 | 0.14521049 | 0.14505224 | 0.10484315 | 0.23441397 |
| **T24_F19** | 0.080878125 | 0.15637189 | 0.22487918 | 0.23632198 | 0.23820434 | 0.20989877 | 0.24593061 | 0.25740657 | 0.15844828 | 0.20241623 | 0.19456689 | 0.18493597 | 0.55844339 | 0.039494063 | 0.03516029 | 0.24803768 | 0.090460932 |
| **T4_F4** | 0.23684389 | 0.25913759 | 0.25495676 | 0.2167992 | 0.45135324 | 0.47476353 | 0.25361324 | 0.63402245 | 0.36635954 | 0.35680801 | 0.27376698 | 0.27377711 | 0.26142467 | 0.060097986 | 0.04818489 | 0.2199271 | 0.14134646 |
| **T8_F4** | 0.23488327 | 0.34000299 | 0.43760096 | 0.42130301 | 0.52653115 | 0.5196866 | 0.44519404 | 0.34023093 | 0.27701809 | 0.2874084 | 0.10329162 | 0.1138761 | 0.44264802 | 0.020357777 | 0.017941495 | 0.13462878 | 0.050569224 |
| **T24_F4** | 0.21333678 | 0.31438511 | 0.3962015 | 0.43403161 | 0.35625657 | 0.35654486 | 0.45947723 | 0.15832664 | 0.20212368 | 0.24336333 | 0.035877408 | 0.043501563 | 0.25315342 | 0.006325899 | 0.00669715 | 0.13554543 | 0.018452381 |
| **T4_PA** | 0.007271727 | 0.007863521 | 0.00719872 | 0.006400996 | 0.014573801 | 0.020670996 | 0.006107237 | 0.04454343 | 0.018577336 | 0.01685546 | 0.17149907 | 0.12217697 | 0.015849363 | 0.19595142 | 0.14881439 | 0.025058275 | 0.1623985 |
| **T8_PA** | 0.031553024 | 0.022597155 | 0.026419678 | 0.021915074 | 0.038950522 | 0.033411669 | 0.023755974 | 0.077625571 | 0.055677408 | 0.045510759 | 0.13468635 | 0.15388669 | 0.040988775 | 0.10218978 | 0.12343548 | 0.040965067 | 0.11502783 |
| **T24_PA** | 0.023692078 | 0.10303005 | 0.11732359 | 0.12512859 | 0.15143663 | 0.12141301 | 0.12806367 | 0.1316922 | 0.072962484 | 0.18386601 | 0.16394558 | 0.19853905 | 0.31904379 | 0.073844565 | 0.064232064 | 0.15259945 | 0.10285445 |
| **T4_RL10** | 0.020900605 | 0.022725765 | 0.022067441 | 0.020747622 | 0.045338147 | 0.055409959 | 0.021623173 | 0.11960422 | 0.05537151 | 0.060255954 | 0.20757444 | 0.20973269 | 0.054869501 | 0.26033058 | 0.19722222 | 0.074000847 | 0.29262926 |
| **T8_RL10** | 0.086476744 | 0.089632852 | 0.084372874 | 0.074003795 | 0.14319985 | 0.16676528 | 0.082030124 | 0.27744367 | 0.16612152 | 0.17155445 | 0.30447052 | 0.30135659 | 0.13464804 | 0.090562844 | 0.077216801 | 0.15412844 | 0.20109976 |
| **T24_RL10** | 0.18472485 | 0.20859328 | 0.22083934 | 0.18458802 | 0.30015178 | 0.27132064 | 0.20708185 | 0.31004663 | 0.24238745 | 0.27537532 | 0.1152812 | 0.1101034 | 0.21992238 | 0.021836541 | 0.023116276 | 0.092018255 | 0.055394485 |

| | T8_F16 | T24_F16 | T4_F16F19 | T8_F16F19 | T24_F16F19 | T4_F19 | T8_F19 | T24_F19 | T4_F4 | T8_F4 | T24_F4 | T4_PA | T8_PA | T24_PA | T4_RL10 | T8_RL10 | T24_RL10 |
|---|---|---|---|---|---|---|---|---|---|---|---|---|---|---|---|---|---|
| **RedCedar** | 0.029446269 | 0.17878928 | 0.004512171 | 0.022584839 | 0.10741389 | 0.007782716 | 0.025543033 | 0.080878125 | 0.23684389 | 0.23488327 | 0.21333678 | 0.007271727 | 0.031553024 | 0.023692078 | 0.020900605 | 0.086476744 | 0.18472485 |
| **T4_RC_Biofilm** | 0.032862191 | 0.2673833 | 0.005641875 | 0.022068966 | 0.11466646 | 0.00764058 | 0.023160583 | 0.15637189 | 0.25913759 | 0.34000299 | 0.31438511 | 0.007863521 | 0.022597155 | 0.10303005 | 0.022725765 | 0.089632852 | 0.20859328 |
| **T8_RC_Biofilm** | 0.033355423 | 0.28539028 | 0.006135794 | 0.022511771 | 0.11866025 | 0.008670197 | 0.026953228 | 0.22487918 | 0.25495676 | 0.43760096 | 0.3962015 | 0.00719872 | 0.026419678 | 0.11732359 | 0.022067441 | 0.084372874 | 0.22083934 |
| **T24_RC_Biofilm** | 0.032040658 | 0.28857627 | 0.004928205 | 0.018050383 | 0.094321242 | 0.006258941 | 0.023790118 | 0.23632198 | 0.2167992 | 0.42130301 | 0.43403161 | 0.006400996 | 0.021915074 | 0.12512859 | 0.020747622 | 0.074003795 | 0.18458802 |
| **T4_C20** | 0.05958267 | 0.32755506 | 0.011181834 | 0.051358147 | 0.1596047 | 0.015847861 | 0.050066644 | 0.23820434 | 0.45135324 | 0.52653115 | 0.35625657 | 0.014573801 | 0.038950522 | 0.15143663 | 0.045338147 | 0.14319985 | 0.30015178 |
| **T8_C20** | 0.07384484 | 0.32103838 | 0.013349515 | 0.047029315 | 0.2005418 | 0.019206497 | 0.058991739 | 0.20989877 | 0.47476353 | 0.5196866 | 0.35654486 | 0.020670996 | 0.033411669 | 0.12141301 | 0.055409959 | 0.16676528 | 0.27132064 |
| **T24_C20** | 0.033498996 | 0.32213641 | 0.005237887 | 0.02149265 | 0.14488253 | 0.006967798 | 0.026098967 | 0.24593061 | 0.25361324 | 0.44519404 | 0.45947723 | 0.006107237 | 0.023755974 | 0.12806367 | 0.021623173 | 0.082030124 | 0.20708185 |
| **T4_D14** | 0.1540792 | 0.29254518 | 0.030134446 | 0.097748262 | 0.25483476 | 0.045626505 | 0.12766411 | 0.25740657 | 0.63402245 | 0.34023093 | 0.15832664 | 0.04454343 | 0.077625571 | 0.1316922 | 0.11960422 | 0.27744367 | 0.31004663 |
| **T8_D14** | 0.084842764 | 0.25195179 | 0.014080196 | 0.05050738 | 0.24019628 | 0.018989653 | 0.060967761 | 0.15844828 | 0.36635954 | 0.27701809 | 0.20212368 | 0.018577336 | 0.055677408 | 0.072962484 | 0.05537151 | 0.16612152 | 0.24238745 |
| **T24_D14** | 0.11082588 | 0.27187786 | 0.0168492 | 0.047382826 | 0.22951889 | 0.021154203 | 0.073258152 | 0.20241623 | 0.35680801 | 0.2874084 | 0.24336333 | 0.01685546 | 0.045510759 | 0.18386601 | 0.060255954 | 0.17155445 | 0.27537532 |
| **T4_F14** | 0.25913242 | 0.087854902 | 0.06366782 | 0.21903052 | 0.13450292 | 0.10653266 | 0.26494789 | 0.19456689 | 0.27376698 | 0.10329162 | 0.035877408 | 0.17149907 | 0.13468635 | 0.16394558 | 0.20757444 | 0.30447052 | 0.1152812 |
| **T8_F14** | 0.25312184 | 0.087523932 | 0.072371222 | 0.21024419 | 0.16976444 | 0.10883797 | 0.30466989 | 0.18493597 | 0.27377711 | 0.1138761 | 0.043501563 | 0.12217697 | 0.15388669 | 0.19853905 | 0.20973269 | 0.30135659 | 0.1101034 |
| **T24_F14** | 0.081991215 | 0.29657956 | 0.010824313 | 0.05901044 | 0.16859519 | 0.016405873 | 0.066060127 | 0.55844339 | 0.26142467 | 0.44264802 | 0.25315342 | 0.015849363 | 0.040988775 | 0.31904379 | 0.054869501 | 0.13464804 | 0.21992238 |
| **T4_F14F16** | 0.26853707 | 0.018485514 | 0.22197055 | 0.18710359 | 0.032809773 | 0.18200409 | 0.14521049 | 0.039494063 | 0.060097986 | 0.020357777 | 0.006325899 | 0.19595142 | 0.10218978 | 0.073844565 | 0.26033058 | 0.090562844 | 0.021836541 |
| **T8_F14F16** | 0.25319865 | 0.016255443 | 0.22043628 | 0.17340426 | 0.037982058 | 0.1573499 | 0.14505224 | 0.03516029 | 0.04818489 | 0.017941495 | 0.00669715 | 0.14881439 | 0.12343548 | 0.064232064 | 0.19722222 | 0.077216801 | 0.023116276 |
| **T24_F14F16** | 0.08195341 | 0.34393064 | 0.023955774 | 0.094136418 | 0.4451631 | 0.022097775 | 0.10484315 | 0.24803768 | 0.2199271 | 0.13462878 | 0.13554543 | 0.025058275 | 0.040965067 | 0.15259945 | 0.074000847 | 0.15412844 | 0.092018255 |
| **T4_F16** | 0.39076758 | 0.051116646 | 0.21777422 | 0.23117803 | 0.067626381 | 0.21130952 | 0.23441397 | 0.090460932 | 0.14134646 | 0.050569224 | 0.018452381 | 0.1623985 | 0.11502783 | 0.10285445 | 0.29262926 | 0.20109976 | 0.055394485 |
| **T8_F16** | 1 | 0.055460542 | 0.17199391 | 0.19543693 | 0.058497673 | 0.15471966 | 0.18743961 | 0.054529894 | 0.1530782 | 0.059869036 | 0.029397645 | 0.13805522 | 0.15 | 0.093401487 | 0.21667552 | 0.19468683 | 0.072868027 |
| **T24_F16** | 0.055460542 | 1 | 0.011526896 | 0.032461027 | 0.27786202 | 0.01206373 | 0.042105263 | 0.283943 | 0.3069528 | 0.34718731 | 0.36188324 | 0.010454396 | 0.019298937 | 0.19135665 | 0.03573614 | 0.117465 | 0.22664885 |
| **T4_F16F19** | 0.17199391 | 0.011526896 | 1 | 0.18022235 | 0.046837851 | 0.33710692 | 0.19917582 | 0.039391937 | 0.029993335 | 0.010827803 | 0.004358814 | 0.11596958 | 0.064305685 | 0.040108401 | 0.1355398 | 0.054589478 | 0.014812923 |
| **T8_F16F19** | 0.19543693 | 0.032461027 | 0.18022235 | 1 | 0.078547925 | 0.20177384 | 0.29533469 | 0.11919757 | 0.10148836 | 0.040312801 | 0.013140009 | 0.1077147 | 0.19714739 | 0.087678229 | 0.21510097 | 0.16498559 | 0.067182533 |
| **T24_F16F19** | 0.058497673 | 0.27786202 | 0.046837851 | 0.078547925 | 1 | 0.033149171 | 0.10231239 | 0.23132814 | 0.22843126 | 0.17833812 | 0.15246937 | 0.023277042 | 0.065124772 | 0.13383617 | 0.065471551 | 0.17299062 | 0.13767434 |
| **T4_F19** | 0.15471966 | 0.01206373 | 0.33710692 | 0.20177384 | 0.033149171 | 1 | 0.34042553 | 0.067624021 | 0.048367594 | 0.015652854 | 0.004996993 | 0.15518745 | 0.087460955 | 0.058124174 | 0.17155425 | 0.076757223 | 0.019546868 |
| **T8_F19** | 0.18743961 | 0.042105263 | 0.19917582 | 0.29533469 | 0.10231239 | 0.34042553 | 1 | 0.14108881 | 0.1338389 | 0.047252747 | 0.017772917 | 0.13938053 | 0.15024121 | 0.15159694 | 0.21728395 | 0.19739574 | 0.061152394 |
| **T24_F19** | 0.054529894 | 0.283943 | 0.039391937 | 0.11919757 | 0.23132814 | 0.067624021 | 0.14108881 | 1 | 0.24768387 | 0.38570325 | 0.19181535 | 0.030061892 | 0.056375541 | 0.49701563 | 0.087533809 | 0.15042076 | 0.22364274 |
| **T4_F4** | 0.1530782 | 0.3069528 | 0.029993335 | 0.10148836 | 0.22843126 | 0.048367594 | 0.1338389 | 0.24768387 | 1 | 0.41543293 | 0.18377306 | 0.047893949 | 0.10700613 | 0.16044029 | 0.10824365 | 0.27356948 | 0.29636121 |
| **T8_F4** | 0.059869036 | 0.34718731 | 0.010827803 | 0.040312801 | 0.17833812 | 0.015652854 | 0.047252747 | 0.38570325 | 0.41543293 | 1 | 0.43511135 | 0.016081871 | 0.034332649 | 0.22090059 | 0.040386827 | 0.13481886 | 0.26324407 |
| **T24_F4** | 0.029397645 | 0.36188324 | 0.004358814 | 0.013140009 | 0.15246937 | 0.004996993 | 0.017772917 | 0.19181535 | 0.18377306 | 0.43511135 | 1 | 0.005795368 | 0.018081073 | 0.097872986 | 0.015423341 | 0.066249415 | 0.15575026 |
| **T4_PA** | 0.13805522 | 0.010454396 | 0.11596958 | 0.1077147 | 0.023277042 | 0.15518745 | 0.13938053 | 0.030061892 | 0.047893949 | 0.016081871 | 0.005795368 | 1 | 0.13610889 | 0.10440376 | 0.18136952 | 0.095607763 | 0.015236668 |
| **T8_PA** | 0.15 | 0.019298937 | 0.064305685 | 0.19714739 | 0.065124772 | 0.087460955 | 0.15024121 | 0.056375541 | 0.10700613 | 0.034332649 | 0.018081073 | 0.13610889 | 1 | 0.088006231 | 0.12375691 | 0.18667557 | 0.091068506 |
| **T24_PA** | 0.093401487 | 0.19135665 | 0.040108401 | 0.087678229 | 0.13383617 | 0.058124174 | 0.15159694 | 0.49701563 | 0.16044029 | 0.22090059 | 0.097872986 | 0.10440376 | 0.088006231 | 1 | 0.10049307 | 0.093455463 | 0.20040746 |
| **T4_RL10** | 0.21667552 | 0.03573614 | 0.1355398 | 0.21510097 | 0.065471551 | 0.17155425 | 0.21728395 | 0.087533809 | 0.10824365 | 0.040386827 | 0.015423341 | 0.18136952 | 0.12375691 | 0.10049307 | 1 | 0.2327856 | 0.060501509 |
| **T8_RL10** | 0.19468683 | 0.117465 | 0.054589478 | 0.16498559 | 0.17299062 | 0.076757223 | 0.19739574 | 0.15042076 | 0.27356948 | 0.13481886 | 0.066249415 | 0.095607763 | 0.18667557 | 0.093455463 | 0.2327856 | 1 | 0.16447065 |
| **T24_RL10** | 0.072868027 | 0.22664885 | 0.014812923 | 0.067182533 | 0.13767434 | 0.019546868 | 0.061152394 | 0.22364274 | 0.29636121 | 0.26324407 | 0.15575026 | 0.015236668 | 0.091068506 | 0.20040746 | 0.060501509 | 0.16447065 | 1 |

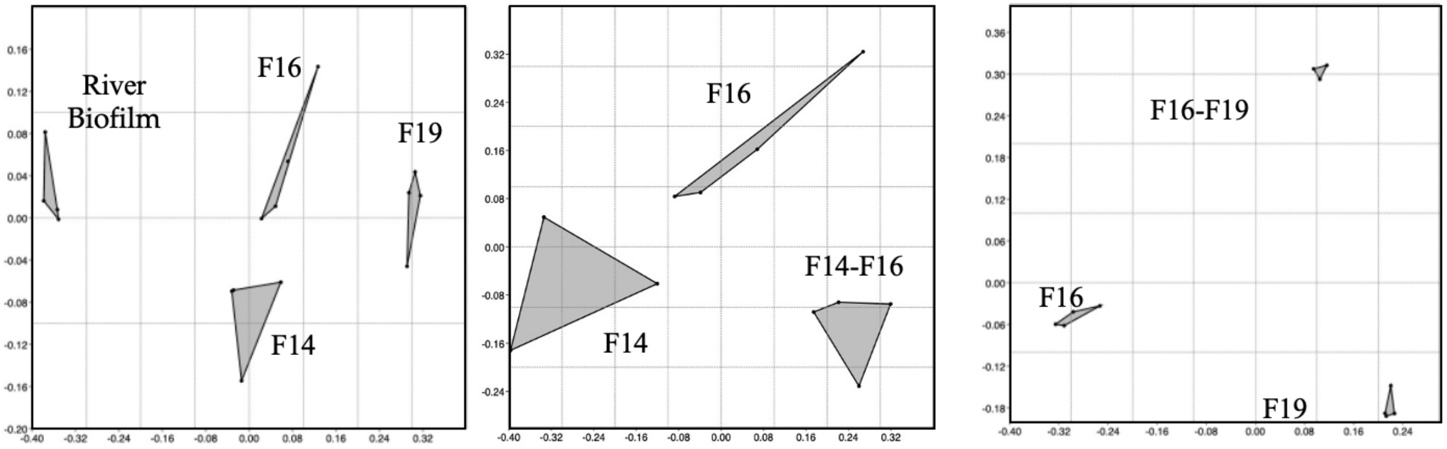


| ANOSIM | River biofilm | F14 | F16 | F19 | F14-F16 | F16-F19 | PERMANOVER | River biofilm | F14 | F16 | F19 | F14-F16 | F16-F19 |
|---|---|---|---|---|---|---|---|---|---|---|---|---|---|
| River biofilm | | 0.027 | 0.026 | 0.028 | 0.0274 | 0.0293 | River biofilm | | 0.028 | 0.028 | 0.029 | 0.0289 | 0.0282 |
| F14 | 1 | | 0.028 | 0.028 | 0.0261 | 0.0298 | F14 | 7.264 | | 0.029 | 0.026 | 0.0296 | 0.0294 |
| F16 | 1 | 0.583 | | 0.027 | 0.0302 | 0.0273 | F16 | 7.245 | 2.549 | | 0.03 | 0.0287 | 0.0296 |
| F19 | 1 | 1 | 1 | | 0.0294 | 0.0269 | F19 | 9.421 | 6.099 | 4.736 | | 0.0314 | 0.0292 |
| F14-F16 | 1 | 1 | 0.813 | 1 | | 0.0291 | F14-F16 | 8.861 | 4.46 | 2.87 | 6.603 | | 0.0282 |
| F16-F19 | 1 | 1 | 0.944 | 1 | 1 | | F16-F19 | 7.796 | 5.458 | 3.776 | 4.239 | 4.602 | |

**Supplemental Fig 5. NMDS, ANOSIM and PERMANOVER analyses of biofilm communities from river and pre-established biofilms of *Hydrogenophaga* F14, *Brevundimonas* F16, and *Acidovorax* F19 at 4 hours incubation**. Founding populations of the pre-established communities have been removed prior to comparative analyses. R values and F values are shaded and uncorrected P values are unshaded.

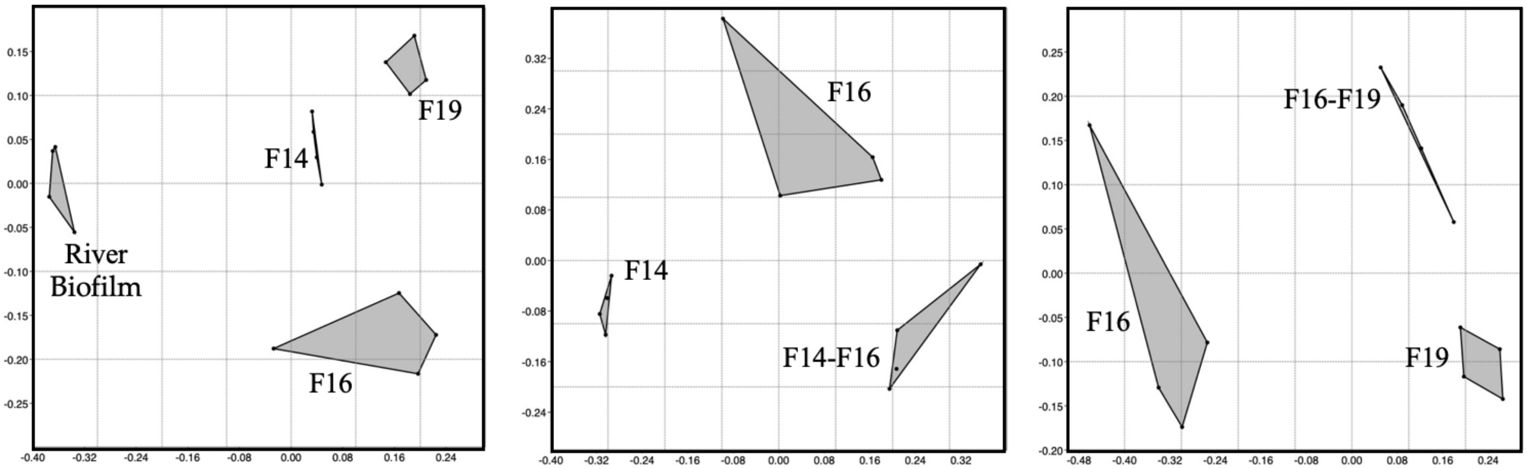


| ANOSIM | River biofilm | F14 | F16 | F19 | F14-F16 | F16-F19 | PERMANOVER | River biofilm | F14 | F16 | F19 | F14-F16 | F16-F19 |
|---|---|---|---|---|---|---|---|---|---|---|---|---|---|
| River biofilm | | 0.031 | 0.026 | 0.03 | 0.0291 | 0.0256 | River biofilm | | 0.029 | 0.028 | 0.03 | 0.027 | 0.0269 |
| F14 | 1 | | 0.028 | 0.03 | 0.0313 | 0.0282 | F14 | 6.047 | | 0.368 | 0.029 | 0.028 | 0.0291 |
| F16 | 1 | 0.906 | | 0.03 | 0.0306 | 0.0294 | F16 | 5.895 | 1.111 | | 0.027 | 0.0298 | 0.0291 |
| F19 | 1 | 1 | 1 | | 0.028 | 0.0293 | F19 | 6.381 | 3.687 | 1.604 | | 0.0264 | 0.0295 |
| F14-F16 | 1 | 1 | 0.677 | 1 | | 0.0303 | F14-F16 | 6.458 | 5.289 | 1.304 | 7.798 | | 0.0285 |
| F16-F19 | 1 | 0.99 | 0.979 | 0.875 | 1 | | F16-F19 | 6.444 | 4.489 | 1.495 | 2.866 | 2.556 | |

**Supplemental Fig 5. NMDS, ANOSIM and PERMANOVER analyses of biofilm communities from river and pre-established biofilms of *Hydrogenophaga* F14, *Brevundimonas* F16, and *Acidovorax* F19 at 8 hours incubation**. Founding populations of the pre-established communities have been removed prior to comparative analyses. R values and F values are shaded and uncorrected P values are unshaded.
